# Mapping Slow Speckle Dynamics to Probe Cellular Metabolic Activity In Vivo using Laser Speckle Contrast Imaging

**DOI:** 10.64898/2026.04.02.713027

**Authors:** Emily Long, Matthew G. Simkulet, Rockwell P. Tang, John Jiang, Şefik Evren Erdener, Timothy M. O’Shea, David A. Boas, Xiaojun Cheng

**Affiliations:** Neurophotonics Center, Boston University, Boston, USA; Institute of Neurological Sciences and Psychiatry, Hacettepe University, Ankara, Turkey

**Keywords:** ischemic stroke, laser speckle contrast imaging

## Abstract

**Significance:** Laser speckle contrast imaging (LSCI) is widely used to measure blood flow, but speckle fluctuations may also encode biologically meaningful dynamics beyond perfusion. Foundational studies in dynamic light scattering (DLS) and micro-optical coherence tomography (μOCT) have also demonstrated that slow coherent signal fluctuations can arise from energy-dependent intracellular motion in *in vitro* and *ex vivo* systems. Building upon these advances, recent work has shown that LSCI has the potential to detect slow speckle dynamics (SSD) correlated with cellular dynamics *in vivo*. However, the biophysical mechanisms underlying SSD in intact brain tissues remain insufficiently validated. Establishing a mechanistic bridge from controlled *ex vivo* and *in vitro* conditions to *in vivo* brain measurements is critical for translating speckle-based imaging beyond perfusion measurements to enable label-free assessment of cellular and metabolic activity in disease models.

**Aim:** The objective of this study is to investigate the biophysical origin of the SSD *in vivo* and evaluate its sensitivity to intracellular metabolic activity in brain tissue.

**Approach:** We utilize an epi-illumination LSCI system to measure speckle contrast as a function of camera exposure time and extract characteristic decorrelation time constants. SSD was investigated in acute mouse brain slices, where blood flow is absent, to eliminate vascular confounds. Cellular metabolism was systematically modulated using 2-deoxyglucose and glucose. Complementary *in vivo* measurements were performed to reveal SSD’s response to hyperoxia and normoxia after ischemic stroke.

**Results:** SSD signals persisted in acute brain slices in the absence of blood flow. Inhibition of glycolysis significantly reduced SSD, while restoration of metabolic substrates partially recovered the signal. In *in vivo* measurements, SSD increased during hyperoxia compared to normoxia after ischemic stroke, suggesting increased oxygen-supported cellular metabolic activity.

**Conclusions:** These results indicate that SSD is sensitive to energy-dependent cellular processes closely tied to metabolic activity. SSD represents a previously uncharacterized, label-free *in vivo* optical contrast that enables assessment of cellular metabolic activity as well as vascular dynamics. This work establishes a mechanistic foundation for using SSD as a general optical marker of cellular viability in *in vivo* measurements.

## 1 Introduction

Laser speckle contrast imaging (LSCI) is a wide-field technique that has been commonly used to monitor cerebral blood flow (CBF)^1–9^. Recently, slow speckle dynamics (SSD) has emerged as a novel contrast mechanism that captures slow speckle temporal fluctuations (∼s) in speckle patterns *in vivo*^10,11^, distinct from conventional LSCI, which is primarily sensitive to fast speckle temporal fluctuations (∼ms) arising from blood flow. In prior work, we demonstrated that SSD can be robustly measured *in vivo* and exhibits strong sensitivity to pathological conditions such as ischemic stroke, revealing spatial patterns that evolve over hours to days following the initiation of multiple stroke models in mice^11^. Unlike the fast speckle dynamics (FSD) that are predominantly driven by the motion of red blood cells, SSD reflects changes occurring on substantially longer timescales, suggesting sensitivity to non-vascular processes within tissue. Importantly, SSD does not require exogenous contrast agents and can be acquired using the same optical system as FSD, making it well-suited for longitudinal measurements. However, the biological origin of the SSD signal remains to be verified.

Clues to the origin of the *in vivo* SSD can be found in prior studies of laser speckle dynamics in *in vitro* and *ex vivo* tissue samples. Pioneering work demonstrated that slow temporal fluctuations in coherent light scattering can arise from intracellular motion, including organelle transport and other metabolically driven processes, and that these dynamics are strongly modulated by ATP availability and cellular viability^12–16^. Consistent with this interpretation, related signals have also been reported using dynamic and micro optical coherence tomography (μOCT), where slow speckle decorrelation has been attributed to active cellular or intracellular motility in living tissues^17^. More specifically, μOCT has enabled direct visualization of ciliary beating, epithelial dynamics, and mucociliary transport in airway tissues, providing striking evidence that long-timescale coherent signal fluctuations can reflect ATP-dependent cellular activity^18,19^. Together, these studies across multiple coherent imaging modalities support the interpretation that optical signals at long timescales can be sensitive to active, energy-dependent cellular processes. However, these mechanistic insights were established using interferometric or depth-resolved optical platforms and have not been directly validated in the context of wide-field LSCI. In particular, it remains unclear whether the biological origins inferred from holographic and μOCT measurements translate to SSD acquired using LSCI in intact brain tissue in vivo, where scattering geometry, spatial averaging, and vascular context differ substantially. Direct experimental confirmation of the contributors to SSD under these conditions is therefore needed.

In this study, we directly investigate the biological contributors to SSD by leveraging the unique ability to apply the same optical measurement framework, epi-illumination LSCI, across both *ex vivo* and *in vivo* preparations. Using acute brain slices, we isolate tissue from confounding vascular flow and systematically perturb metabolic activity, enabling direct assessment of how SSD responds to changes in cellular viability and energy state. We then extend these findings to an *in vivo* ischemic stroke model, examining how SSD evolves in response to oxygenation. By combining controlled *ex vivo* experiments with *in vivo* stroke imaging, our approach bridges a critical gap between mechanistic understanding and physiological relevance. Together, these experiments demonstrate that SSD reflects metabolically relevant tissue dynamics driven by active cellular processes, supporting its interpretation as a functional biomarker of tissue viability across experimental contexts and disease states.

## 2 Materials and Methods

### 2.1 Imaging Setup

#### 2.1.1 Laser Speckle Contrast Imaging

A custom-built epi-illumination LSCI system was used in the study (Fig 1a)^11^. A laser light source (Topica WS, 853 nm) is mounted on an arm perpendicular to the imaging axis. After expansion, the coherent laser light passes through a linear polarizer and a polarizing beam splitter (Thorlabs BSF10-B) before being delivered to the sample. The backscattered light is collected by a 1.5x lens system, passes through a crossed linear polarizer, and is imaged onto a CMOS camera (Basler acA 1440-220umNIR). An iris is placed in front of the camera to adjust the amount of light entering the camera. A custom Basler pylon-based software is used for acquisition and real-time monitoring of speckle contrast. For measuring slow speckle dynamics (SSD), a long camera exposure time of *T* = 100 ms was used with a camera frame rate of 10 Hz. The laser power is adjusted to avoid camera saturation while ensuring the signal is high enough such that camera noise is negligible.

**Fig 1.**
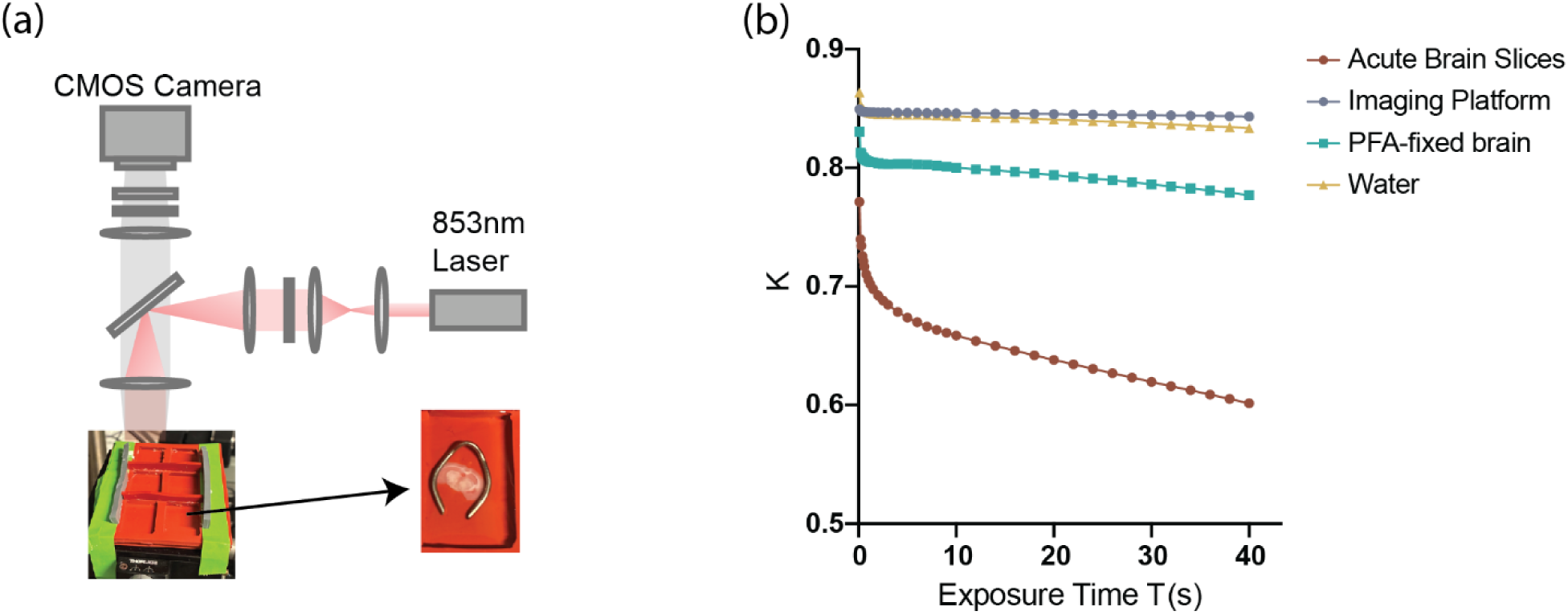
*Ex vivo* validation of slow speckle dynamics (SSD) signal. (a) Schematic of the custom epi-illumination laser speckle contrast imaging (LSCI) system used for *ex vivo* measurements, showing the custom silicone imaging platform and slice harp used to secure brain slices and ensure mechanical stability during imaging. (b) Representative speckle contrast *K* as a function of camera exposure time *T* for acute brain slices, the imaging platform, PFA-fixed brain tissue, and water.

The spatial contrast *K* at a synthetic camera exposure time *T* is calculated as^1,20,21^:

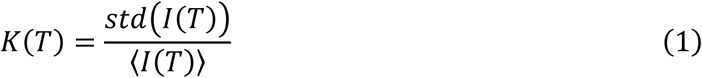

where *std* stands for the standard deviation of the intensity *I* and 〈⋯ 〉 represents the average over a set of pixels. The contrast *K* was calculated for each pixel over a window of 7×7 pixels and frames were averaged over the entire acquisition period. We measured spatial contrast at the long exposure time *T* = 100 ms for 240 s to get a sufficient number of frames to average.

#### 2.1.2 Optical Coherence Tomography

Optical coherence tomography (OCT) imaging was performed to quantify tissue scattering changes during stroke progression. OCT signal attenuation was computed from the depth-dependent decay of the OCT intensity signal^22^. Imaging was conducted using a spectral domain OCT system (1310 nm center wavelength, bandwidth 170 nm, Thorlabs) with a 5× objective lens (Mitutoyo), following previously reported acquisition parameters^23^. Volumetric OCT data were acquired over a field of view of 3.5 mm × 3.5 mm with 1200 × 1200 pixels, corresponding to a voxel size of ∼3 µm × 3 µm × 3.5 µm. For each time point, three volumetric scans were acquired and averaged for subsequent analysis to improve signal-to-noise ratio.

OCT attenuation maps were computed using custom MATLAB code^23^. The signal intensity was calculated as the logarithm of the averaged signal, and the attenuation coefficient at each lateral pixel was determined as the slope of a first-order linear fit to the depth-resolved intensity profile. Fits were performed over a depth range of ∼ 120 µm, beginning approximately 250–300 µm below the cranial window to avoid surface artifacts.

### 2.2 Ex Vivo Measurements

#### 2.2.1 Brain Slice Preparation

Acute coronal brain slices were prepared from adult C57BL/6J mice (12–16 weeks old; 3 males and 3 females). Mice were given an overdose of isoflurane (4% at 0.7L/min) and immediately transcardially perfused with 30 mL of ice-cold artificial cerebrospinal fluid (aCSF; in mM: 124 NaCl, 3 KCl, 1.25 NaH₂PO₄, 26 NaHCO₃, 2 CaCl₂, 1 MgSO₄, and 10 glucose) to remove residual blood, thereby reducing hemoglobin absorption and minimizing vascular contributions to LSCI signals. Brains were rapidly extracted and sectioned into 300-µm-thick coronal slices using a Compresstome (Precisionary Instruments, VF-300-OZ). Approximately 25 coronal slices were obtained per brain, from which 4–6 slices at the hippocampal level were selected for subsequent measurements. Selected slices were incubated in oxygen-equilibrated aCSF at ∼4°C for 30 min. The aCSF was oxygen-equilibrated overnight by placing the solution in an oxygen incubator prior to use.

Slices were then transferred to a custom-cut silicone imaging platform designed with individual recessed wells (∼2 cm × 3 cm × 1 cm) such that each slice was positioned in a separate compartment that is manually shaped to fit the desired recording area (Fig 1a). Silicone is selected for its chemical inertness and non-absorbent properties, ensuring that treatment compounds remain at stable concentrations during the experiment, as well as for its ability to provide reliable tissue adherence. Each slice was immersed in room-temperature aCSF and gently secured with a dedicated slice harp (Warner Instruments, SHD-26H/15) to prevent motion during recovery and imaging. Slices were allowed to equilibrate for approximately 1 h prior to recording.

The aCSF solution used in the experiment was pre-equilibrated overnight in a cell culture incubator (37 °C, 5% CO₂) prior to use. pH was monitored using a calibrated pH meter during experiments and confirmed to remain stable at ∼7.4 across measurements, minimizing potential confounding effects from pH variation.

#### 2.2.2 Experimental Design

For metabolic perturbation experiments, baseline imaging was first acquired in standard aCSF. The bath solution was then manually exchanged with aCSF containing one of the following perturbations: 2-deoxy-D-glucose (2-DG; 10 mM) to inhibit glycolysis, or elevated glucose (final concentration 30 mM) to increase substrate availability for glycolytic metabolism. Following solution exchange, slices were incubated under the new condition for 15–20 min prior to imaging to allow physiological effects of the perturbation to develop. At the conclusion of each experiment, slices were exposed to 4% paraformaldehyde (PFA) in phosphate-buffered saline (PBS) to induce metabolic arrest (Fig 2a). This terminal condition served as a negative control to confirm signal sensitivity and to distinguish treatment-driven changes from imaging drift. For data acquisition, each SSD measurement lasted 4 min.

**Fig 2.**
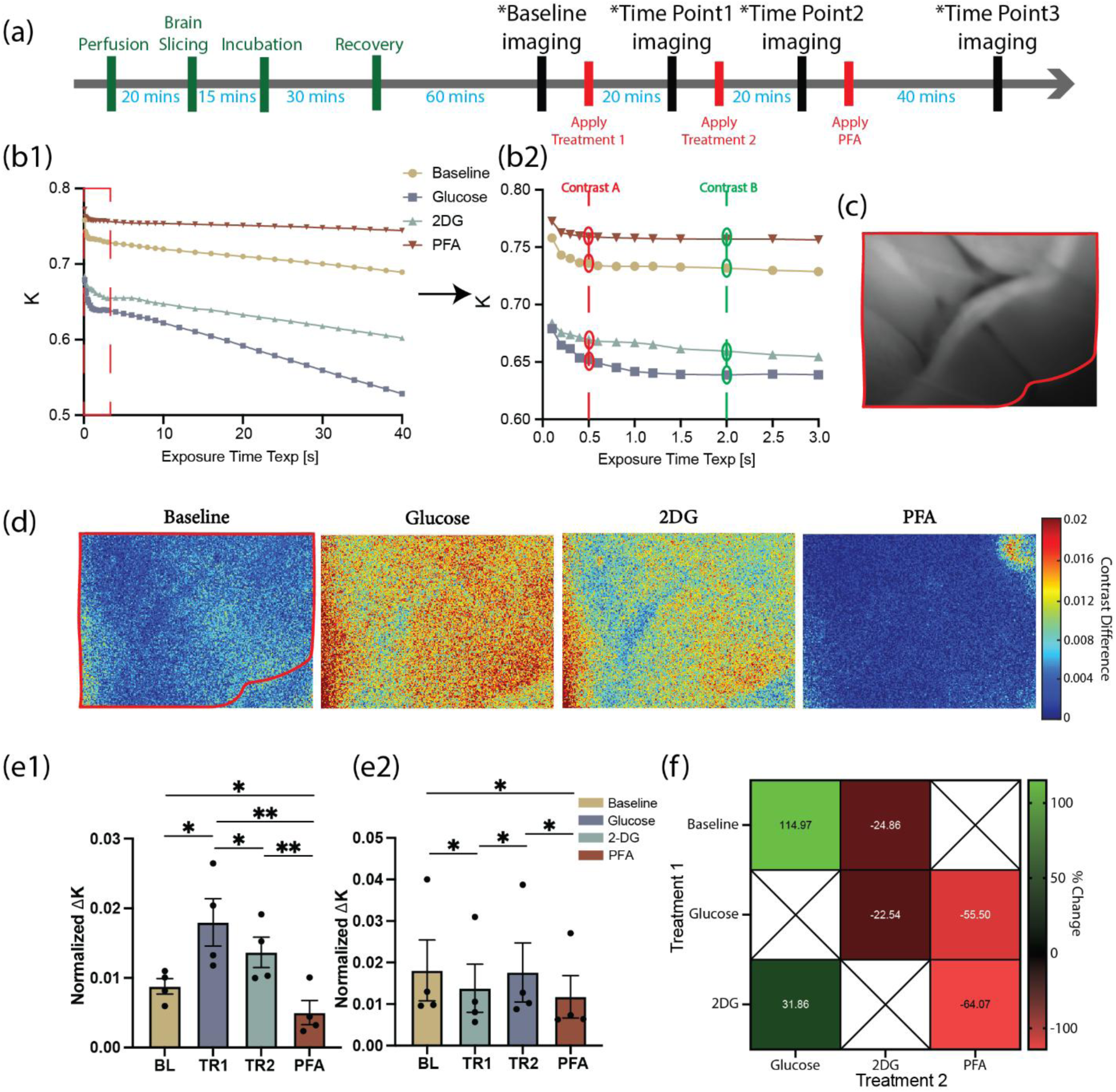
*Ex vivo* slow speckle dynamics (SSD) measurements in acute brain slices under metabolic perturbations. (a) Experimental timeline for *ex vivo* measurements. (b1) Representative speckle contrast curves acquired under baseline and metabolic perturbation conditions. (b2) Zoomed-in view of selected exposure times used to compute the SSD contrast difference metric, Δ*K*. (c) Acute brain slice viewed under the widefield camera with selected region of interest (ROI) outlined for Δ*K* analysis. (d) Representative spatial maps of Δ*K* under different metabolic conditions. (e1-e2) Quantification of normalized Δ*K* across experimental conditions. Data are shown as mean ± SEM with individual data points overlaid; statistical significance is indicated (\**p* < 0.05), performed by one-tailed t-test. (f) Heatmap summarizing percent changes in Δ*K* for all pairwise combinations of sequential treatments. % change is calculated as 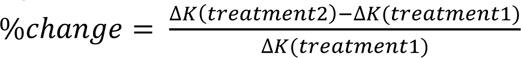

All *ex vivo* experiments were conducted in room-temperature aCSF without continuous carbogen bubbling during imaging. Continuous bubbling was not used because the SSD and FSD measurements are highly sensitive to mechanical motion, and bubble-induced surface disturbances introduce substantial speckle artifacts. Slice viability under these conditions was supported by the preservation of dynamic signals throughout the experiment and by the robust signal suppression observed following PFA fixation. All *ex vivo* measurements were completed within 6 h of animal euthanasia.

### 2.3 In Vivo Measurements

#### 2.3.1 Animal Preparation

All animal procedures were approved by the Boston University Institutional Animal Care and Use Committee and conducted in accordance with the National Institutes of Health Guide for the Care and Use of Laboratory Animals. Adult C57BL/6J mice (12–16 weeks old; 3 males and 5 females) were used in this study and housed under standard light–dark conditions with ad libitum access to food and water.

For optical imaging, a cranial window was implanted over the right frontoparietal cortex. Following established protocols^24,25^, a ∼4-mm diameter craniotomy was performed and sealed with a glass coverslip (Warner Instruments). A custom aluminum head bar was superglued to the skull to enable head fixation during imaging. Mice were allowed to recover for at least two weeks following surgery before imaging experiments were conducted.

#### 2.3.2 Stroke Model

Focal cerebral ischemia was induced using the distal middle cerebral artery (dMCA) FeCl₃ thrombosis model following established protocols^26^. Prior to stroke induction, a ∼2-mm diameter craniotomy was created in the temporal bone to expose the dMCA while preserving the dura. A ferric chloride (FeCl₃) solution (30% w/v in water) was freshly prepared, and a small piece of filter paper (approximately 1 mm × 0.5 mm) was soaked in the solution for 30 s. The filter paper was then placed directly on the dura overlying the dMCA to initiate thrombotic occlusion (Fig 3b). Cerebral blood flow (CBF) was continuously monitored in real time using LSCI to confirm the onset and stability of ischemia. A successful occlusion was defined as a reduction of greater than 50% in CBF relative to baseline, accompanied by the loss of at least one visible pial arterial branch in the LSCI maps. The FeCl₃-soaked filter paper was removed approximately 10 min after placement, and the exposed dura was washed and kept hydrated with sterile saline. After the last hyperoxia measurement (Section 2.3.3), the cranial opening was sealed with bone wax and dental cement prior to subsequent imaging measurements.

**Fig 3.**
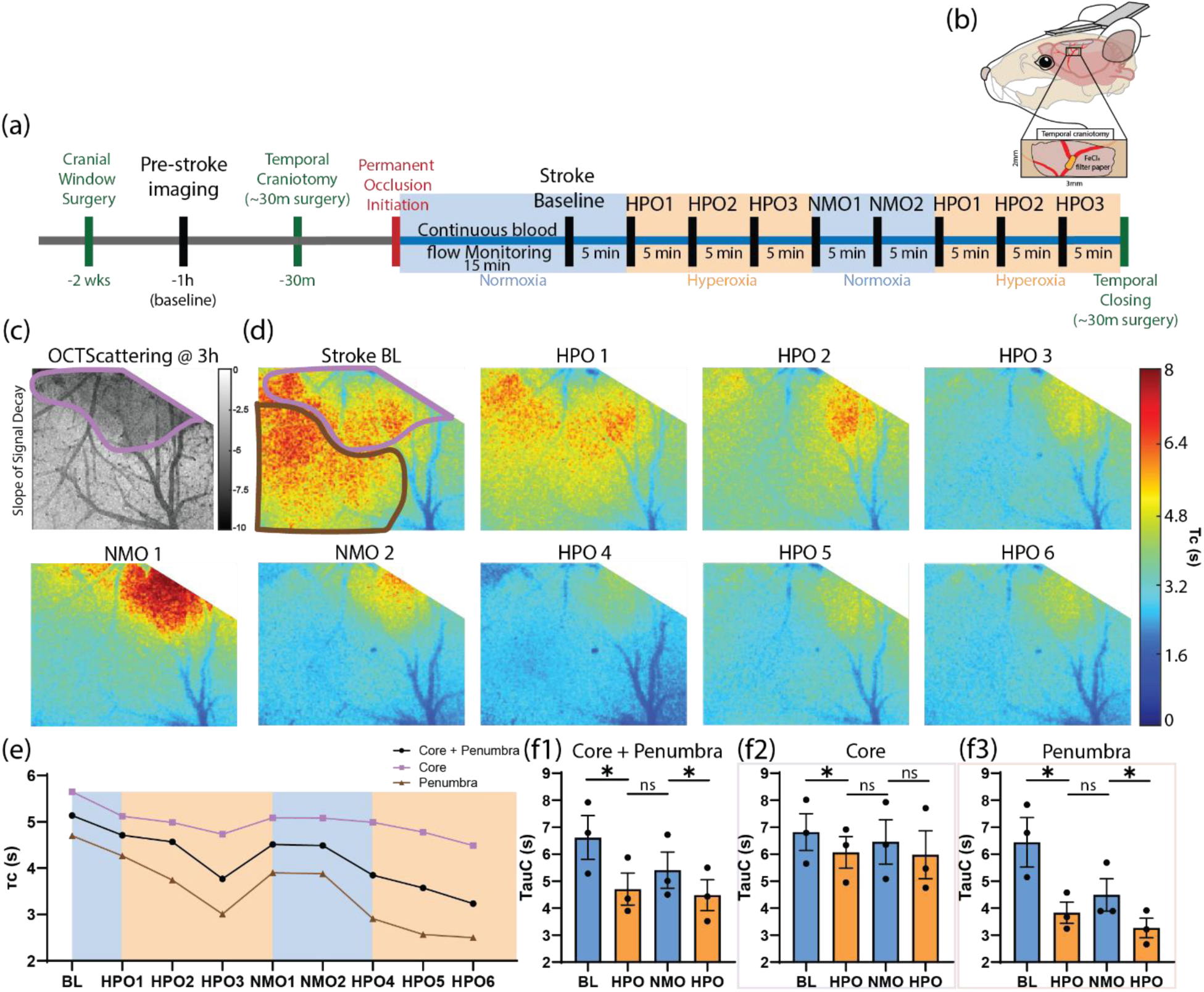
*In vivo* slow speckle dynamics (SSD) reveal differential metabolic responses to hyperoxia and normoxia after ischemic stroke. (a) Experimental timeline for in vivo measurements. (b) Schematic of permanent FeCl₃-induced middle cerebral artery occlusion model. (c) OCT scattering map at 3 h post-stroke, with the stroke core outlined in purple and used to define core. (d) Representative spatial maps of the SSD time constant (*τ*_*C*_) at stroke baseline (BL), successive hyperoxia (HPO1–HPO3), normoxia (NMO1–NMO2) blocks, and subsequent hyperoxia (HPO4-6) blocks with stroke core outlined in purple while penumbra outlined in brown. (e) Representative time course of *τ*_*C*_ averaged over core, penumbra, and combined regions. (f1–f3) Quantification of *τ*_*C*_ within the combined core + penumbra region (f1), stroke core only (f2), and penumbra only (f3). Statistical comparisons were performed using paired tests across conditions (*p < 0.05; ns, not significant.

#### 2.3.3 Experimental Design

All imaging sessions were performed under isoflurane anesthesia (3% for induction, 1% for maintenance and during imaging). Baseline measurements (4 min SSD) were obtained prior to induction of the stroke.

Oxygen modulation began 15 min after stroke initiation while mice remained anesthetized and spontaneously breathing through a nose cone. Normoxic and hyperoxic conditions were delivered by switching the inlet gas supply between room air and 100% O₂. Gas switching was performed at the outlet manifold immediately upstream of the nose cone to minimize dead volume and ensure rapid changes in the inspired gas composition.

During each gas condition, 4 min of SSD imaging was acquired repeatedly. Prior to each SSD acquisition, real-time monitoring was reviewed to confirm the absence of spreading depolarization events; acquisitions coinciding with spreading depolarizations were excluded from analysis. Mice underwent three consecutive measurements under hyperoxia, followed by two consecutive measurements under room air, and then three additional measurements under hyperoxia (Fig 3a). Throughout all imaging sessions, body temperature was continuously monitored and maintained at 37℃ using a feedback-controlled heating pad (Kent Scientific) to ensure physiological stability. Temperature control was employed to minimize potential confounding effects of temperature fluctuations on speckle dynamics measurements.

### 2.4 Data Analysis

For *in vivo* measurements, the decorrelation time *τ*_*c*2_ of SSD was quantified by fitting the decay curve of the spatial speckle contrast *K*(*T*) with a nonlinear model. To reduce parameter crosstalk between fast and slow dynamics, SSD fitting was performed independently of FSD by using only long exposure times, following previously reported approaches^11^. Under this condition, the spatial *K* is dominated by SSD contributions and can be described by the SSD model when using a long exposure time *T* ≥100 ms:

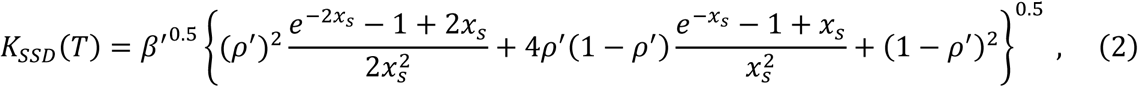

with 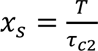. The parameter 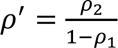 is derived from 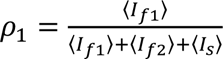 and *ρ*_2_ = 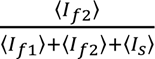, which represent the fraction of dynamically scattered light contributing to FSD and SSD, respectively; *I*_*f*1_, *I*_*f*2_, and *I*_*s*_ are the fast-fluctuating, slow-fluctuating, and static portions of the scattered light intensities, respectively. The parameter *β*^′^ = *β*(1 − *ρ*_1_)^2^ is derived from *ρ*_1_ and *β*, which accounts for coherence loss due to speckle averaging, polarization effects, and system instability.

SSD fitting was performed for each pixel using synthetic exposure times ranging from 1 s to 40 s. Model fitting was carried out using custom MATLAB scripts with constrained nonlinear least-squares optimization. *β*^′^ is approximated during fitting by constraining it to the *K*_*SSD*_^2^ calculated at the first timepoint on the decay curve. *ρ*^′^ was constrained between 0 and 1 in fitting. SSD is quantified by *τ*_*c*2_, where a larger *τ*_*c*2_ value arises from slower cellular dynamics.

For *ex vivo* measurements, SSD data were analyzed using the contrast difference metric (ΔK) instead. The nonlinear decorrelation model discussed above was not applicable here because the contrast curve *ex vivo* exhibited multi-component behavior, inconsistent with the assumption of the model (see Discussion). Δ*K* was defined as the pixelwise normalized difference in speckle contrast between long (2 s) and short (0.5 s) exposure times:

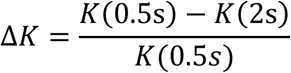

Here a larger Δ*K* represents faster signal decorrelation and therefore faster dynamics. The time points 0.5 s and 2 s are chosen based on the time scale of the slow dynamics (∼1 s), which we found to be sensitive to relevant physiological perturbations rather than other artifacts such as water surface vibrations. All data analysis was performed using custom MATLAB scripts (MathWorks) and GraphPad Prism. Unless otherwise stated, data are reported as mean ± standard error of the mean (SEM). Paired statistical comparisons were performed when measurements were obtained from the same animal or slice across conditions. Statistical tests and sample sizes are specified in the corresponding figure captions, with statistical significance defined as p < 0.05.

## 3 Results

### 3.1 Slow speckle dynamics (SSD) sensitivity to metabolic perturbation in acute brain slices

To assess whether slow speckle dynamics (SSD) reflect intrinsic tissue activity rather than static optical properties, speckle contrast measurements were compared between acute brain slices and non-dynamics controls in *ex vivo* preparations (Fig. 1a). Compared to controls, including the silicone imaging platform, water, and paraformaldehyde (PFA)-fixed tissue, acute brain slices exhibited a distinct speckle contrast profile over exposure time *T* (Fig. 1b). While non-dynamic samples showed minimal variation across exposure times, acute brain tissues displayed pronounced decay behavior, indicating the presence of endogenous biological dynamics. The marked separation between living tissue and static or non-living controls demonstrates that SSD arises from intrinsic tissue activity and remains sensitive to physiological processes in the absence of blood flow.

To further decipher the contributors to the SSD signal, metabolic perturbations were introduced to test its sensitivity to changes in cellular metabolic state (Fig. 2a). Glucose serves as the primary substrate for glycolysis and supports adenosine triphosphate (ATP) production in brain tissue^27,28^, whereas 2-deoxy-D-glucose (2DG; Sigma-Aldrich, 25972-M), a glucose analog, is phosphorylated by hexokinase but cannot proceed through glycolysis, thereby inhibiting glycolytic flux and reducing cellular energy availability^29–31^. 4% Paraformaldehyde (PFA) fixation was used as a terminal condition to induce metabolic arrest and confirm tissue viability prior to fixation. Consistent with these biochemical mechanisms, representative speckle contrast curves exhibited systematic changes following metabolic manipulation (Fig. 2b1). Inhibition of glycolysis with 2DG was associated with a clear reduction in the rate of decay in *K*, whereas increased glucose availability produced a faster decay in contrast, reflecting opposing effects on SSD dynamics. PFA fixation resulted in near-complete loss of contrast decay, further confirming the dependence of the signal on viable tissue activity. To quantify these effects, a normalized SSD analysis metric, Δ*K*, was introduced as previously mentioned in the methods section. Δ*K* was defined as the difference between speckle contrast values at 0.5 s and 2 s exposure times, normalized by *K*(0.5s). This exposure-time pair was selected empirically because the resulting contrast difference consistently showed the greatest sensitivity to metabolic perturbations (Fig. 2b2). Decorrelation on the order of ∼ 1s also aligns with previously reported timescales of intracellular energetic and cytoskeletal processes observed in dynamic speckle fluctuation studies of living tissue^32^. Notably, an additional component was observed at longer exposure times (approximately 3–30 s); however, this component exhibited minimal sensitivity to metabolic manipulation which could arise from slow vibrations of the brain slices. Using this normalized metric, spatial distributions of Δ*K* were examined across acute brain slices under different metabolic conditions. Regions of interest (ROIs) were selected from the camera view to avoid tissue edges and areas exhibiting visible slicing artifacts or structural disruption (Fig. 2c). Representative Δ*K* maps revealed clear modulation of SSD signals following metabolic perturbation, with distinct spatial patterns observed under glucose supplementation, glycolytic inhibition with 2DG, and terminal PFA fixation (Fig. 2d). In particular, 2DG treatment consistently produced reduced Δ*K* values across the selected regions, whereas increased glucose availability resulted in relative increases in Δ*K* values. PFA fixation yielded near-zero Δ*K*, indicating loss of dynamic tissue activity. To assess reproducibility, group-level analysis was performed in 8 brain slices from 6 mice. To account for potential order effects, two treatment sequences were employed swapping the order of treatments administered (Fig. 2e1-e2). Across both sequences, Δ*K* demonstrated consistent decreases following glycolytic inhibition and strong suppression following fixation, while glucose supplementation produced relative increases toward baseline levels. To further examine transition-dependent effects, percent changes in Δ*K* were computed relative to the preceding condition and summarized as a transition heatmap (Fig. 2f). In this representation, the y-axis denotes the initial treatment and the x-axis denotes the subsequent treatment. The heatmap highlights systematic reductions in SSD following transitions to metabolic inhibition and increases following transitions to enhanced substrate availability, independent of treatment order. Overall, these findings indicate that SSD provides a robust, metabolism-sensitive optical readout of intrinsic tissue activity in acute brain slices, independent of vascular contributions.

### 3.2 In vivo SSD reveals oxygen-dependent modulation of metabolic dynamics following ischemic stroke

To evaluate the sensitivity of SSD-derived metrics to physiologically relevant perturbations *in vivo*, we performed longitudinal imaging during alternating hyperoxia and normoxia. An ischemic stroke model was used (Fig. 3b) to enhance signal sensitivity to oxygen modulation relative to healthy cortex. SSD imaging was initiated 15 min after stroke induction and acquired continuously during alternating hyperoxia and normoxia experimental blocks (Fig. 3a–b). For in vivo experiments, SSD dynamics were quantified using the fitted speckle decorrelation time constant (*τ*_*C*_). At stroke baseline (15 min post-initiation), elevated *τ*_*C*_ values were observed within the ischemic territory. During hyperoxia blocks, *τ*_*C*_ decreased across large portions of the imaged cortex, whereas normoxia was associated with stabilization or partial recovery of *τ*_*C*_ (Fig. 3d). This pattern was reproducible across repeated oxygenation cycles, indicating that SSD signals dynamically track acute changes in oxygenation in the ischemic brain.

To enable regional analysis, optical coherence tomography (OCT) scattering imaging was performed at 3h post-stroke to define the stroke core (Fig. 3c). Spatial inspection revealed that the amplitude of *τ*_*C*_modulation varied across regions, suggesting heterogeneous SSD sensitivity within ischemic tissue (Fig. 3d). Quantitative comparison across conditions confirmed a significant reduction in *τ*_*C*_ during hyperoxia relative to baseline in the combined core and penumbra region (Fig. 3f1). When analyzed separately, *τ*_*C*_ changes within the stroke core were modest and did not differ significantly between hyperoxia and normoxia (Fig. 3f2). In contrast, the penumbra exhibited a more apparent decrease in *τ*_*C*_during hyperoxia compared to both baseline and normoxia (Fig. 3f3), indicating enhanced SSD activity in peri-infarct regions.

Together, these results demonstrate that SSD-derived time constants are sensitive to acute oxygen perturbations *in vivo* in a stroke disease model, with region-dependent responses between core and penumbra. This experiment establishes the dynamic responsiveness of SSD signals under physiologically relevant modulation and provides a foundation for subsequent analyses of SSD as a functional marker of tissue metabolic states.

## 4 Discussion

In this study, we demonstrate slow speckle dynamics (SSD) as an optical readout of intrinsic tissue metabolic state, both *ex vivo* in acute mouse brain slices and *in vivo* following ischemic stroke. In *ex vivo* preparations, SSD signals clearly distinguished living brain tissue from static or non-dynamic controls and responded systematically to targeted metabolic perturbations, including glycolytic inhibition, substrate enhancement, and terminal paraformaldehyde (PFA) fixation. These results establish that SSD reflects active, viability-dependent tissue dynamics independent of blood flow dynamics. Extending these findings *in vivo*, we show that SSD-derived time constants dynamically respond to acute oxygenation changes after ischemic stroke, with more pronounced modulation in the penumbra relative to the infarct core. Together, these experiments demonstrate that SSD captures metabolically relevant tissue dynamics across experimental contexts and disease states, supporting its potential as a functional biomarker.

Notably, time constant fitting established in prior work^11^ was not applied to the *ex vivo* experiments because the existing fitting framework was developed specifically for *in vivo* measurements, where SSD decay curves are well captured by a single effective time constant within the exposure-time range typically analyzed (0-40 s). In contrast, analysis of *ex vivo* speckle contrast as a function of exposure time revealed multiple regimes of temporal behavior, indicating that the single-component fitting model developed *in vivo* does not adequately capture the underlying dynamics in this context. While a single exposure-time pair (0.5–2 s) was selected for quantitative analysis due to its consistent sensitivity to metabolic perturbations, additional components were observed at shorter (0–0.5 s) and longer (3–30 s) exposure ranges. Comparison with non-biological controls revealed that the short-timescale component closely resembled the behavior observed in a sample that only contains water, suggesting that it is dominated by nonbiological effects such as water evaporation rather than intrinsic tissue activity. In contrast, the long-timescale component exhibited minimal sensitivity to metabolic manipulation and was not reliably suppressed by fixation, suggesting that it does not reflect active cellular dynamics but could arise from slice vibrations etc. Progressive tissue swelling was observed in some slices toward the end of the experimental timeline, consistent with osmotic imbalance or gradual metabolic deterioration, and may contribute to this long-timescale behavior through altered scattering and reduced intracellular mobility. Together, these findings indicate that the long-timescale component likely reflects slow cumulative structural or hydration equilibration rather than active cellular dynamics. These observations motivated the use of Δ*K* in the intermediate regime, while avoiding application of a fitting model whose underlying assumptions are not satisfied in the *ex vivo* setting.

*In vivo*, alternating hyperoxia and normoxia revealed rapid and reversible modulation of SSD-derived time constants following stroke. Hyperoxia was associated with reduced *τ*_*C*_values, particularly in the penumbra, while normoxia was associated with increased or unchanged *τ*_*C*_values. These effects likely reflect oxygen-dependent regulation of mitochondrial metabolism and cellular activity in peri-infarct tissue, where metabolic function is impaired but not abolished. The muted response observed in the ischemic core is consistent with severe metabolic failure and limited residual cellular activity, whereas the penumbra retains sufficient viability to exhibit dynamic metabolic modulation^33,34^. Notably, the directionality of SSD changes observed *in vivo* is consistent with the responses measured *ex vivo*. Whether metabolic suppression arises from ischemic injury or chemical inhibition with 2-deoxy-D-glucose (2DG), both conditions are associated with reduced SSD activity, reflected as increased *τ*_*C*_ *in vivo* and decreased Δ*K ex vivo*. Conversely, oxygen supplementation in the stroke model produces SSD changes consistent with enhanced metabolic activity, analogous to the effects of glucose supplementation observed in acute slices. This concordance further supports a shared metabolic interpretation of SSD across experimental contexts and indicates that *τ*_*C*_ and Δ*K* provide internally consistent quantifications of slow speckle dynamics.

Previous literature has established the use of speckle dynamics to probe intracellular motion driven by active biological processes rather than blood flow^12–15,35,36^, with long-exposure speckle fluctuations persisting in avascular systems and being strongly suppressed by metabolic inhibition via administration of iodoacetate or temperature manipulation. Within this framework, SSD is understood to reflect slow intracellular and subcellular motions, such as organelle transport, cytoskeletal remodeling, and membrane fluctuations, activities that depend on cellular energy availability. Perturbations to glycolytic flux therefore provide a direct means of modulating these processes, offering a mechanistic explanation for the graded SSD responses observed under metabolic inhibition, substrate supplementation, and fixation. The present *ex vivo* experiments provide controlled validation of this metabolic sensitivity in the absence of perfusion, while the *in vivo* oxygenation experiments using the same LSCI set-up extend this interpretation to a pathophysiologically relevant setting, where oxygen availability dynamically regulates residual metabolic activity in ischemic tissue.

Importantly, although coherent light scattering and micro–optical coherence tomography (μOCT) studies have elegantly demonstrated ATP-dependent speckle dynamics at the cellular scale^12–19^, these modalities rely on interferometric phase sensitivity and micron-scale resolution that render them susceptible to bulk motion and mechanical instability *in vivo*. In phase-sensitive OCT systems, nanometer- to micron-scale displacements can introduce substantial decorrelation or phase noise, complicating interpretation of slow temporal fluctuations^37^. Such sensitivity poses practical challenges in in vivo brain measurements, where respiration, cardiac pulsation, and subtle drift of the cranial window introduce unavoidable motion artifacts^38^. In contrast, LSCI employs wide-field spatial ensemble averaging and intensity-based contrast analysis. By sacrificing microscopic spatial resolution in exchange for improved statistical robustness, LSCI mitigates sensitivity to physiological motion and enables reliable detection of slow speckle dynamics under realistic *in vivo* conditions, including ischemic injury. Thus, the present work translates mechanistically grounded speckle measurements into a framework compatible with longitudinal imaging in intact brain disease models.

Several experimental considerations warrant discussion. In *ex vivo* slice experiments, while aCSF was oxygen-equilibrated prior to imaging, continuous carbogen bubbling was not employed during acquisition. Active bubbling introduces fluid motion and mechanical perturbations that are incompatible with SSD measurements, which are highly sensitive to subtle tissue and surface dynamics; intermittent bubbling between imaging blocks can further disrupt tissue positioning and stability, introducing substantial variability. As a result, oxygen delivery relied on passive diffusion, which may introduce gradients across slice thickness and contribute to temporal drift in metabolic state. Throughout all experiments, however, aCSF pH was continuously monitored and remained stable, helping to mitigate confounding effects related to acid–base imbalance. The absence of measurable acidification in the bathing medium is also consistent with maintained tissue viability and suggests that severe oxygen deprivation, which would be expected to induce lactic acidosis and proton efflux from damaged cells^39^, did not occur during the imaging period.

In addition, the near-complete suppression of SSD signals following paraformaldehyde fixation at the end of each protocol confirms that the tissue remained viable and metabolically active up to the experimental endpoint. Although treatment comparisons were internally consistent and reproducible across slices, future implementations incorporating motion-isolated or perfusion-stabilized oxygen delivery could further improve metabolic control while preserving measurement stability. In addition, variability in Δ*K* was observed across acute brain slices, even under nominally identical experimental conditions. Such variability is expected in *ex vivo* preparations and may arise from differences in slice thickness, baseline metabolic state, and the extent of mechanical stress incurred during slicing and handling. While these effects introduce variability in absolute Δ*K* values across slices, the directionality and relative changes induced by metabolic perturbations were consistent, supporting the robustness of SSD as a metabolism-sensitive metric when interpreted in a within-slice or transition-based framework.

Looking forward, the metabolic sensitivity of slow speckle dynamics suggests broad applicability across disease models characterized by altered cellular energetics, including ischemic stroke, cancer, and neurodegenerative disorders. Prior work^40^ has demonstrated the utility of slow speckle dynamics for probing intracellular motility and testing cellular response to anticancer treatment in *ex vivo* cancer models, supporting the potential application of this approach to disease models. Taken together, this study establishes SSD as a label-free, metabolism-sensitive optical readout that is independent of blood flow and responsive to physiologically relevant perturbations across *ex vivo* and *in vivo* settings. Importantly, the present demonstration of SSD using LSCI extends foundational work on speckle-based cellular dynamics imaging^12–16^ and dynamic micro–optical coherence tomography^17–19^ from *in vitro* and *ex vivo* preparations to *in vivo* disease models. By providing access to intrinsic tissue dynamics linked to cellular energy availability with computational simplicity and scalability, SSD complements existing imaging modalities and holds promise as an *in vivo* functional biomarker for assessing tissue viability and metabolic state in diseases.

## Disclosures

The authors declare no conflicts of interest.

## Acknowledgments

This work was supported by the National Institutes of Health (R01NS127156, T32NS136080, UG3EB034710). We thank Bingxue Liu, Dmitry Postnov, and Jianbo Tang for building and maintaining the imaging systems used in this study, with special recognition to Bingxue Liu for laying the foundation of the project. We also thank Alexander Howard, Allen Zhou, John Giblin, and Xin Brown for helpful discussions. This study was facilitated by the Boston University Neurophotonics Center.

## Notes

### Competing Interest Statement

The authors have declared no competing interest.

